# High contrast, moving targets in an emerging target paradigm promote fast visuomotor responses during visually guided reaching

**DOI:** 10.1101/2021.01.27.428509

**Authors:** Rebecca A. Kozak, Brian D. Corneil

**Affiliations:** Graduate Program in Neuroscience, Western University, London, On, Canada, N6A 5B7; Department of Psychology, Western University, London, Ontario, Canada, N6A 5B7; Department of Physiology and Pharmacology, Western University, London, Ontario, Canada, N6A 5B7; Robarts Research Institute, 1151 Richmond St. N, London, Ontario, Canada, N6A 5B7

**Author notes:** Corresponding Author: Brian D. Corneil.

**Keywords:** visually-guided reaches, EMG, SLR, moving targets, contrast, reaction time, RT

## Abstract

Humans have a remarkable capacity to rapidly interact with the surrounding environment, often by transforming visual input into motor output on a moment-to-moment basis. But what visual features promote rapid reaching? High contrast, fast-moving targets elicit strong responses in the superior colliculus (SC), a structure associated with express saccades and implicated in rapid electromyographic (EMG) responses on upper limb muscles. To test the influence of stimulus properties on rapid reaches, we had human subjects perform visually guided reaches to moving targets varied by speed (experiment 1) or speed and contrast (experiment 2), in an emerging target paradigm which has recently been shown to robustly elicit fast visuomotor responses. Our analysis focused on stimulus-locked responses (SLRs) on upper limb muscles. SLRs appear within <100 ms of target presentation, and as the first wave of muscle recruitment, they have been hypothesized to arise from the SC. Across 32 subjects studied in both experiments, 97% expressed SLRs in the emerging target paradigm, whereas only 69% expressed SLRs in an immediate response paradigm towards static targets. Faster moving targets (experiment 1) evoked large magnitude SLRs, while high contrast fast moving targets (experiment 2) evoked short latency, large magnitude SLRs. In some instances, SLR magnitude exceeded the magnitude of movement aligned activity. Both large magnitude and short latency SLRs were correlated with short latency reach reaction times. Our results support the hypothesis that, in scenarios requiring expedited responses, a subcortical pathway originating in the SC elicits the earliest wave of muscle recruitment, expediting reaction times.

**New & Noteworthy:** How does the brain rapidly transform vision into action? Here, by recording upper limb muscle activity, we find that high contrast and fast moving targets are highly effective at evoking rapid visually guided reaches. We surmise that a brainstem circuit originating in the superior colliculus contributes to the most rapid reaching responses. When time is of the essence, cortical areas may serve to prime this circuit, and elaborate subsequent phases of recruitment.

## Introduction

To perform a visually-guided reach, for example to pick-up a baseball, visual information such as the shape and size of the ball is transformed into motor commands which are communicated to the motor periphery. However, if the ball is thrown to us, time is of the essence and we must perform a rapid reach in order to catch the ball. In such situations, the latency of the response depends on the visual attributes of the stimulus; for example, earlier responses are elicited by larger size, higher contrast, or lower spatial frequency targets (Veerman et al., 2008; Wood et al., 2015; Kozak et al., 2019). Other work has suggested that the most rapid visuomotor transformation for reaching is mediated by a tecto-reticulo-spinal pathway, which lies in parallel with the better studied corticospinal pathway (Day and Lyon, 2000; Pruszynski et al., 2010; Gu et al., 2016; Glover and Baker, 2019).

To better understand the most rapid visuomotor transformations for limb control, we and others have recorded electromyographic (EMG) activity from upper limb muscles during rapid visually-guided reaches. Doing so has led to the identification of stimulus-locked responses (SLRs) which are the first wave of muscle activity influenced by the visual stimulus (SLRs are also termed rapid visuomotor responses by (Glover and Baker, 2019) or express responses by (Contemori et al., 2021)). SLRs evolve within < 100 ms of stimulus onset, which is before some response latencies in visual and motor cortices (Maunsell and Gibson, 1992; Schmolesky et al., 1998; Lara et al., 2018), but after response latencies in the superior colliculus (SC) from which the tecto-reticulo-spinal track originates (Stuphorn et al., 1999; Rezvani and Corneil, 2008). Further, responses in the SC and SLRs exhibit similar stimulus preferences, in that both are preferentially evoked by high contrast (Li and Basso, 2008; Marino et al., 2012; Wood et al., 2015) or low spatial frequency (Chen et al., 2018; Kozak et al., 2019) stimuli. Such observations are consistent with the hypothesis that the SC mediates the SLR.

Robust SC responses are also elicited by moving visual targets (Marrocco and Li, 1977), however, a systematic investigation of target motion on the SLR has not been performed. Previous reports examining short latency reaching responses have typically relied on targets that either suddenly appear (a *static target paradigm*; (Pruszynski et al., 2010)), or are displaced during a reach (*target jump paradigm*; (Soechting and Lacquaniti, 1983; Day and Lyon, 2000)). Recently, we have presented an *emerging target paradigm*, where a target travels down an inverted ‘Y’ path before disappearing and then emerging from behind an occluder (Kozak et al., 2020). Subjects are instructed to reach and intercept the target when it emerges from behind the occluder. Benefits of the emerging target paradigm include the stability of the arm when the target emerges (which is not the case in the target jump paradigm), and the generation of robust fast visuomotor responses in almost every subject (which is not the case in the static target paradigm). In Kozak et al. (2020), all subjects exhibited at SLR in the emerging target paradigm with magnitudes five times larger than in the static paradigm. The efficacy of the emerging target paradigm in generating SLRs has been replicated in a larger sample of subjects, wherein 20 of 21 subjects expressed an SLR (Contemori et al., 2021). Thus, essentially every subject has the circuitry to generate fast visuomotor responses; the failure to do so in past studies arose from the reliance on suboptimal stimuli and/or tasks.

The main goal of the current study is to systematically investigate how target speed and contrast impact fast visuomotor responses in the emerging target paradigm. We hypothesize that the SC mediates rapid visuomotor responses on the upper limb, and therefore predict that high contrast fast moving targets will elicit more prevalent, shorter latency, and larger magnitude SLRs. A secondary goal is to better understand how the emerging target paradigm promotes the generation of SLRs, by comparing such SLRs to those in the static target paradigm. Overall, we find that fast moving, high-contrast, targets in the emerging target paradigm robustly recruits the circuitry mediating the most rapid visuomotor transformations for reaching movements.

## Materials and methods

### Subjects

A total of 34 subjects (18 females, 16 males; mean age: 24.2 years SD: 5.6) completed at least one of the two experiments. Two subjects completed both experiments. All subjects provided informed consent, were paid for their participation, and were free to withdraw from the experiment at any time. All but 1 subject was right-handed. All subjects had normal or corrected-to-normal vision, with no current visual, neurological, or musculoskeletal disorders. All procedures were approved by the Health Science Research Ethics Board at the University of Western Ontario. Two subjects (1 from each experiment) were excluded due to low quality EMG recordings (see below).

### Apparatus

Subjects were seated in either a KINARM robotic Exoskeleton (experiment 1) or a KINARM End-Point Lab (experiment 2) (KINARM Technologies, Kingston, ON, Canada), and performed reaching movements with their right arm. Both platforms relied on the same experimental software. Visual stimuli were projected onto an upward facing mirror from a downward facing monitor in experiment 1 (LG 47LS35A; size: 47” resolution: 1920x1080 pixels (Exoskeleton)) or a custom built in projector in experiment 2 (PROPixx projector by VPixx, Saint-Bruno, QC, Canada; custom integrated into the KINARM End-Point Lab). Each platform has a different distance from the eyes to the target. We positioned visual stimuli at different locations in each platform so that targets were presented at similar retinal eccentricities and velocities. In both platforms, a shield below the mirror occluded direct vision of the hand, but hand position was represented on the monitor in real-time via a *real-time cursor* (RTC) projected onto the screen. A photodiode was used to indicate the precise time of visual stimulus presentation, and all kinematic and electromyographic (EMG) data were aligned to photodiode onset. A loading force was provided in both experiments to increase the baseline activity of the muscle of interest, so that the SLR could be expressed as either an increase or decrease in muscle activity following leftward or rightward target presentation, respectively. This loading force was different in Experiment 1 and 2 given that these experiments were conducted on different platforms.

### Experimental design

In experiment 1 and 2, we used an emerging target paradigm as previously described in ((Kozak et al., 2020) see **Figure 1**). The emerging target paradigm involves a target starting at the top of an inverted y (**Figure 1a**), moving towards an occluder and disappearing behind it (**Figure 1b**), then emerging below the occluder at one of two outlets. Subjects were instructed to reach and intercept the target after it emerged from behind the occlude as quickly and accurately as possible (**Figure 1c**) (Kozak et al., 2020). During target occlusion, subjects were instructed to fixate on a notch at the bottom of the occluder, midway between the inverted Y outlets. The target only became visible once it had fully moved past the occluder (i.e., the target was presented in its entirety, rather than sliding past the occluder to avoid a half-moon stimulus). The target then continued to move down the inverted ‘Y’ path. The trial ended as soon as the RTC contacted the peripheral target, or if the target passed the bottom of the screen. In both experiments, we also collected data in a *static task*, where subjects performed visually guided reaches as quickly and accurately as possible towards stationary targets in a separate block of trials. Doing so allowed us to directly compare SLRs in the emerging target paradigm to those obtained with a static task typically used to evoke the SLR (Pruszynski et al., 2010; Wood et al., 2015; Gu et al., 2016; Kozak et al., 2019). Kinematic and electromyographic (EMG) data are aligned to target emergence below the occluder in the emerging target task, or relative to peripheral target presentation in the static task. Eye movements were not measured.

**Figure 1.**
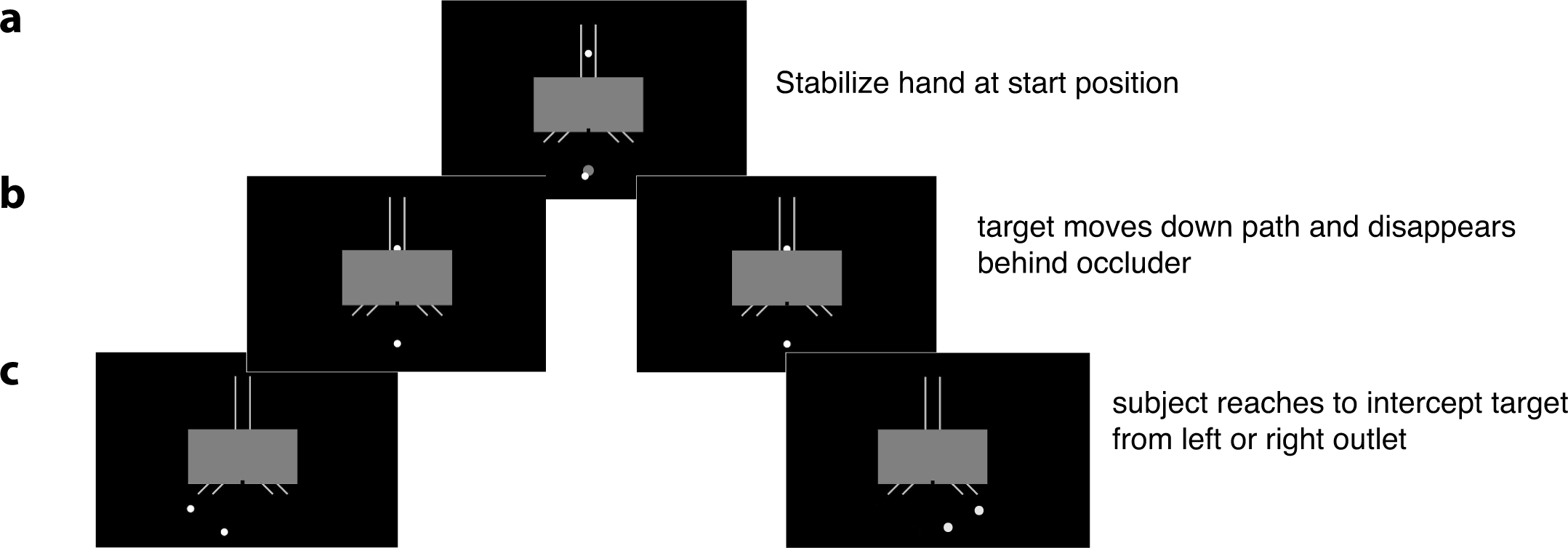
The emerging target paradigm (Kozak et al., 2020). a) A target (T1) appears above an occluder (grey box) and the subject moves the real-time cursor (RTC; white dot below occluder that represents hand position), into the start position (T0; grey dot below the occluder). b) Once the RTC is in the start position, T1 moves towards the subject along the inverted y path and then disappears behind the occluder. c) The subject makes a visually-guided reach to T1 as soon as it emerges at the right or left outlet below the occluder.

### Experiment 1: effects of speed

#### Static task

In the static task in Experiment 1, subjects (*n* = 16) performed visually guided reaches from a stationary arm position to static targets appearing pseudorandomly to either the right or left of the start position. Throughout this task, a constant torque of 2 Nm was applied to the shoulder, increasing tonic activity on the muscle of interest. Each trial was initiated when a subject moved the RTC representing their hand into a white target representing the start position (T0). T0 was 1.5 cm in diameter, located approximately 45 cm in front of the subject relative to midline, and changed to grey once the RTC was in T0. A target (T1) was presented after a 1s hold period. If the subject exited the start position before completion of this hold period, the trial was reset, and T0 changed back to white. Subjects were instructed to also look at T0 before the onset of the target (T1). After the hold period, a white T1 was presented on the black background 17 cm to the left or right of the start position (∼17.5 degrees of visual angle). A 17 cm eccentricity was chosen as it resembled the interception point of the emerging target paradigm. The trial ended when the RTC made contact with the center of T1. T0 remained visible as a grey target, changing back to white once the trial ended. Subjects completed 1 block of 220 trials (110 trials towards each target), before they performed the emerging target paradigm.

#### Emerging target paradigm

After completing the static task, the same 16 subjects performed visually guided reaches towards moving targets in the emerging target paradigm (See **Figure 1** for protocol). The start position, RTC, hold period, background loading torque, and target radius were the same as in the static task.

We implemented three target speeds in our paradigm (10, 15 or 20 cm/s), translating to initial velocities of approximately 10, 15 or 20 retinal degrees per second, respectively. These speeds were chosen to span a range from ‘slow’ to ‘fast’ while still permitting target interception of the fastest target. To control for the time that a participant has to prepare for an upcoming reach, the duration that T1 was visible above the occluder was constant (.5s) regardless of target speed. As targets travelled at different speeds, this meant that they started at different heights above the occluder. The occluder was 30 cm wide by 15 cm high, and the center of the occluder was located ∼63 cm away from the subject. Behind the occluder, the target moved at a constant speed of (15 cm/s) regardless of initial or final target speed. Thus, the target consistently disappeared behind the occluder for 1s on all trials.

We also included a static 0cm/s target within the emerging target paradigm, where a target began moving at a speed of 20 cm/s, disappeared behind the occluder, and then appeared ∼ 17 cm to the left or right of the start position and did not move. Thus, the initial conditions where the target drops into the occluder were the same for the 0 cm/s and 20 cm/s targets. On the 0 cm/s trial, the trial ended when the subjects made contact with the target. The target disappeared behind the occluder for 900 ms on 0 cm/s trials, rather than 1000 ms as on moving target trials, due to an oversight. Therefore, a target appeared below the occluder 1.5 s after initiation for moving targets, and 1.4 s for static targets. Contemori and colleagues (2021) recently showed that SLRs are promoted by certainty of the time of target appearance, which would have predicted that weaker SLRs would have been observed for 0 cm/s targets. This indicates that our results would be biased against robust SLRs in 0cm/s conditions, which we did not observe.

All targets in experiment 1 were white (110 cd/m^2^) presented against a black (.6 cd/m^2^) background (contrast ratio: 183:1). There were 8 unique trial conditions (4 target speeds 0, 10, 15 and 20 cm/s presented to the left and right). Subjects completed 2 blocks of 480 trials, with each block containing 60 repetitions of each unique trial condition presented pseudorandomly without replacement, yielding a total of 120 trials for each unique trial condition.

### Experiment 2: effects of contrast and speed

Experiment 2 was designed to investigate the interaction between target speed and contrast on the SLR. This experiment was conducted in a different apparatus than experiment 1, in aKINARM End-Point system. As in experiment 1, we ran a static task followed by an emerging target paradigm.

#### Static task

In the *static task* in experiment 2, subjects (*n* = 18) performed visually guided reaches initiated from a stationary position, towards static left or right targets. As in experiment 1, each trial was initiated when a subject moved the RTC representing hand position into a start position (T0). Due to different apparatuses being used in experiment 1 and 2, we note relevant differences below. A constant torque was applied through the handle throughout the experiment (2Nm down, 5Nm right) to increase the baseline activity of the muscle of interest. T0 was aligned with the subject’s midline (0, -42 cm relative to the midpoint between the robot manipulanda; radius 1 cm). T0 changed from white to grey once the RTC was moved into it. The trial was reset if the subject exited T0 before completion of the hold period (randomized time between 1000 to 2000 ms). Our use of a randomized hold time in experiment 2 resembles that used in the work by Wood and colleagues (2015), which also manipulated stimulus contrast. Subjects were instructed to foveate T0 before presentation of the target (T1). After the hold period, T1 (target with a 1 cm radius) appeared 10 cm to the left or right (∼15.4 degrees of visual angle). The trial ended as soon as RTC made contact with the peripheral target. There were 4 unique trial conditions; a high contrast white (275 cd/m^2^: contrast ratio 793.8:1) or low contrast grey (18 cd/m^2^: contrast ratio 50:1) target presented against a black (.36 cd/m^2^) background to either the left or right side. T0 remained visible as a grey target, changing back to white once the trial ended. Subjects completed 1 block of 400 pseudorandomized trials, yielding a total of 100 trials for each unique trial condition.

#### Emerging target paradigm

After completing the static task, the same 18 subjects completed the emerging target paradigm. As in experiment 1, subjects performed visually guided reaches initiated from a stable posture towards moving targets that emerged suddenly beneath an occluder. Throughout the experiment, the same torque was applied as used in the static task. In experiment 2, we used a two-by-two design, where we modulated speed (including a ‘slow’ and a ‘fast’ target (10 cm/s, 20 cm/s; translating to approximately 10 and 20 degrees of visual angle); and contrast (the ‘low’ and ‘high’ contrasts). In experiment 2, the occluder was 30cm by 15cm, centered at the midpoint between the manipulanda (0, - 29 cm). As in experiment 1, we controlled for the amount of time that T1 was visible before being obscured by the occluder (.5s). The time T1 disappeared behind the occluder in experiment 2 is shorter (0.5 s) than in experiment 1 (1 s). There were 8 unique trial conditions (2 speeds, 2 contrasts, leftward or rightward targets). The same contrasts were used from the static task. Subjects completed 2 blocks of 400 pseudorandomized trials, yielding a total of 100 trials for each unique trial condition.

#### Data acquisition and analysis

Surface electromyographic (EMG) activity was recorded from the right pectoralis muscle with double-differential surface electrodes (Delsys Inc. Bagnoli-8 system, Boston, MA USA). We placed one surface electrode on the sternal portion of the muscle, and another surface electrode on the clavicular head of the muscle. Further details regarding electrode placement can be found in our previous methods paper (Kozak et al., 2020). Our rationale for placing two electrodes is as a backup, given the potential of losing the adhesion of surface electrodes during a long experiment. Off-line, we analyzed the EMG recording with the higher signal to noise ratio. EMG signals were sampled at 1 kHz, amplified by 1000, full-wave rectified off-line, and smoothed with a 7-point smoothing function. We excluded two subjects with poor quality muscle recordings, which was defined as an increase in peak EMG activity above baseline of less than 40 μV during leftward reaching movements.

Kinematic data were sampled at 1 kHz by the Kinarm data acquisition system in both experiments. Reaction time (RT) was calculated as time from sudden target onset in the static target task, or target emergence below the occluder in the emerging target paradigm (both measured by the photodiode) to reach initiation. Reach initiation in both experiments was defined as 8% of the peak tangential velocity. We excluded trials with RTs less than 120 ms due to presumed anticipation, and also excluded trials with RTs exceeding 500 ms due to presumed inattentiveness. Overall we retained more than 93% of all trials.

As described previously (Kozak et al., 2020), we used a receiver-operating characteristic (ROC) analysis to define the presence and latency of the SLR. Briefly, the time-series ROC analysis indicates the likelihood of discriminating stimulus presentation based on EMG alone. As the SLR is the first wave of muscle activity influenced by the visual stimulus, we rely on this method to discriminate the moment a visual stimulus influences EMG activity. The ROC value (i.e., the area under the ROC curve) was calculated and plotted at every time sample (1ms) from 100 ms before to 300 ms after target presentation (an example may be seen in **Figure 2d** overlaid with mean +/- SE of EMG activity for the data shown in **Figure 2a**). A ROC value of .5 indicates chance discrimination, whereas values of 1 or 0 indicate perfectly correct or incorrect discrimination relative to target presentation, respectively. Discrimination thresholds were set to 0.6, and the discrimination latency was determined as the first of 8 of 10 consecutive points that exceeded this value.

**Figure 2.**
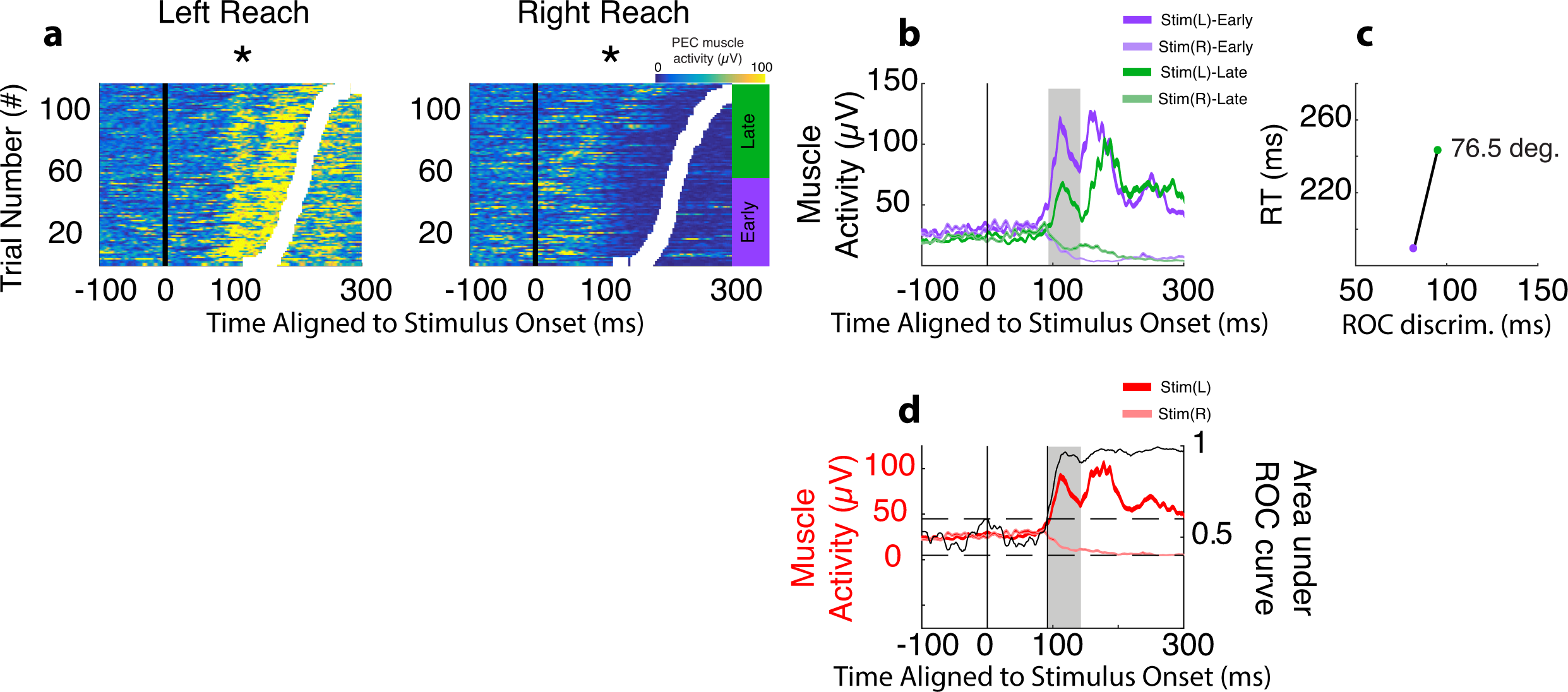
Detection of SLRs. a) Exemplar trial-by-trial recruitment of activity from right pectoralis muscle during leftward and rightward reaches from a subject with a detectable SLR. Each row is a different trial, with the intensity of color conveying degree of recruitment. Trials sorted by RT (white boxes) and aligned to stimulus onset. The SLR (highlighted by asterisk) appears as a vertical banding of increased or decreased recruitment aligned to left or right stimulus presentation, respectively, rather than movement onset. b) Mean +/- SE of EMG activity for the early- (purple) and late- (green) RT subsets based on mean RT. c) In the RT split analysis, separate time-series ROCs were run for the early and late subsets, and then the mean RT for each subset is plotted as a function of the discrimination time. The slope of the line connecting these points determines whether the divergence of EMG activity is more time-locked to stimulus or movement onset. d) For SLR+ observations, all trials were pooled together regardless of RT to determine the SLR latency (derived from the time-series ROC) and magnitude (derived from the mean EMG waveforms; see methods). Grey rectangle in b) and d) depict the SLR interval.

To define the presence of an SLR, an RT split analysis was used (as described in: (Wood et al., 2015; Kozak et al., 2020)). The purpose of the RT split analysis is to empirically test whether EMG activity is more likely to evolve with stimulus presentation, or arm movement over a large number of trials. This method is used as an additional conservative approach to relying on pooled EMG, which may be unfairly biased towards a subset of the shortest latency trials. To perform the RT split analysis, separate time series ROC analyses were conducted on early and late RT trials (purple or green highlighted trials in **Figure 2a**; see also mean EMG traces in **Figure 2b**), defined relative to mean RT. We then plotted the mean RT for the early or late RT subsets versus the ROC discrimination time, and then connected these two points with a line (**Figure 2c**). The slope of this line is used to determine whether the initial change in EMG activity is more locked to stimulus onset (slope ≥ 67.5 deg.) or not (slope < 67.5 deg.). The example shown in **Figure 2** illustrates an example where an SLR was detected (termed SLR+; versus SLR-).

We used the same criterion for determining latency and magnitude of the SLR as used in (Kozak et al., 2020). For SLR+ observations, SLR latency was defined as the first point where the time-series ROC constructed from all trials (regardless of RT) surpassed .6 for 8 out of 10 trials (see **Figure 2d)**. SLR magnitude was defined as the area between leftward and rightward mean EMG traces for an interval spanning from SLR latency to 30 ms later. When comparing SLR magnitudes between subjects, all SLR+ values were first normalized to the maximum within-subject SLR magnitude. For example, if a subject exhibited SLRs in 5 conditions, the largest SLR condition for that subject is given a value of 1 and all other magnitudes are presented as a fraction of 1. When comparing SLR magnitudes to recruitment associated with voluntary movement, we divided the magnitudes attained from the SLR epoch by the sum of the movement-aligned activity with a 30 ms window surrounding peak EMG activity.

### Statistical analyses

Statistical analyses were performed in MATLAB (version R2016a, The MathWorks, Inc., Natick, Massachusetts, United States). When comparing SLR prevalence across conditions, chi-squared tests were used to determine differences in the proportion of subjects exhibiting an SLR.

To compare SLR latencies and magnitudes, as well as reaction times across conditions, we used a one-way ANOVA for experiment 1 and a 2 by 2 ANOVA for experiment 2. Post-hoc pairwise comparisons relied on the Tukey-Kramer critical value, as well as paired and unpaired t-tests when appropriate. Correlational analyses were performed on latency, magnitude and RT measures on a subject-by-subject basis.

## Results

Previous work has shown that the emerging target paradigm is highly effective at eliciting rapid visuomotor responses, such as stimulus-locked responses (SLRs) and short-latency reach reaction times (RTs) (Kozak et al., 2020; Contemori et al., 2021). Here, we sought to investigate the features of the stimulus which best elicit SLRs by altering the speed (experiment 1) or the speed and contrast (experiment 2) of the target, and compared these results to static tasks. We first confirmed that all 32 subjects exhibited an SLR in at least one condition in both experiment 1 and 2 in either the static task or emerging target paradigm (**Figure 3;** each subject had at least one diamond above the dotted line, indicating a detectable SLR). Significantly more subjects exhibited an SLR in the emerging target paradigm compared to the static target task (31 (97%) vs 22 (69%) SLR+ subjects, respectively; chi-squared test: p= 0.0189, χ2= 5.51, df= 1). Our low SLR detection rates in the static task is consistent with previous reports (Wood et al., 2015; Gu et al., 2016; Pruszynski et al., 2018).

**Figure 3.**
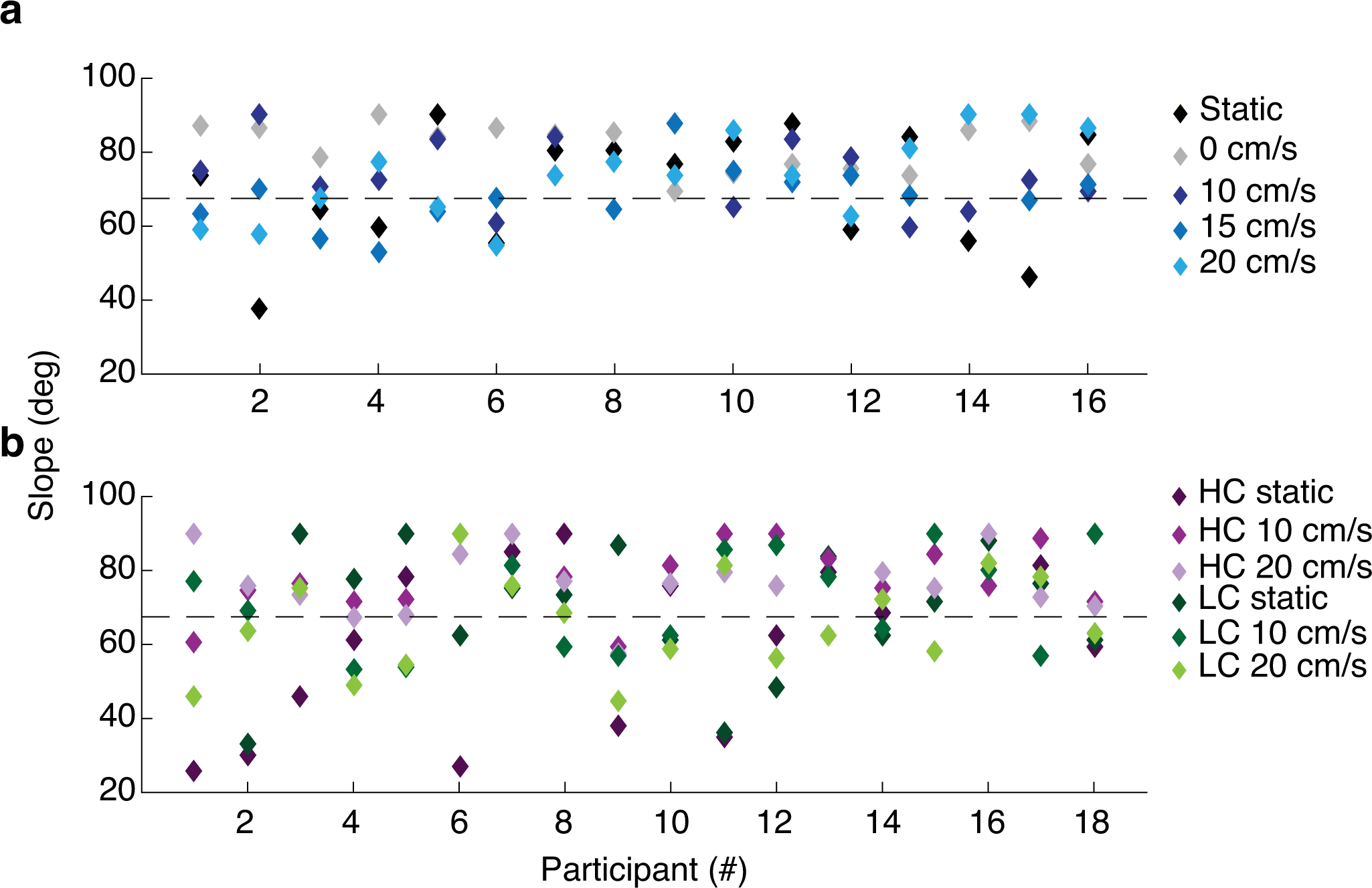
Prevalence of SLRs across the population. Slope of the relationship between discrimination time and average RT for early and late RT groups (e.g. see slope in Figure 2c) Each data point represents a unique subject for a particular condition. Slopes are capped at 90 degrees and a dotted black line is drawn at 67.5 deg. A data point above this line represents a SLR+ observation, whereas a data point below the line represents SLR- observations. a) Experiment 1. Black and grey diamonds represent static conditions in the static and emerging target paradigms, respectively. Blue diamonds represent moving targets in emerging target paradigm, with slower moving targets are represented by darker shades. b) Experiment 2. Purple diamonds represent high contrast targets; green diamonds represent lower contrast targets. Static and slower moving targets are represented by darker shades. 2 subjects completed both experiments; subjects 2 and 16 from experiment 1 are subjects 18 and 12 from experiment 2, respectively.

### Experiment 1: Faster moving targets elicit larger magnitude SLRs

In experiment 1, we investigated whether target speed influences the SLR in an emerging target paradigm. Data from a representative subject is shown in **Figure 4**. In the plots depicting single- trial EMG activity, the SLR appears as a vertical band of EMG activity due to changes occurring around the same time after target emergence on every trial (grey boxes, two left columns in **Figure 4**). The SLR differs from the subsequent phases of EMG activity tied to movement onset (indicated by white boxes demarcating sorted reaction time). **Figure 4** also shows the mean EMG traces, time-series ROC plots, and a depiction of how an SLR is detected (see methods- RT split analysis; slopes >67.5 indicate an SLR is present). In this subject, the SLR began ∼90 ms after target emergence in all conditions, regardless of target speed, or whether a stationary stimulus was presented in the emerging target paradigm or static task. Further, while SLR latency did not appear to change between conditions, it appears that SLR magnitude increased for faster moving targets (e.g., compare the intensity of yellows for the 0 vs 20 cm/s targets in the left column of **Figure 4**, or the peak of mean EMG activity in the SLR interval in the 3^rd^ column).

**Figure 4.**
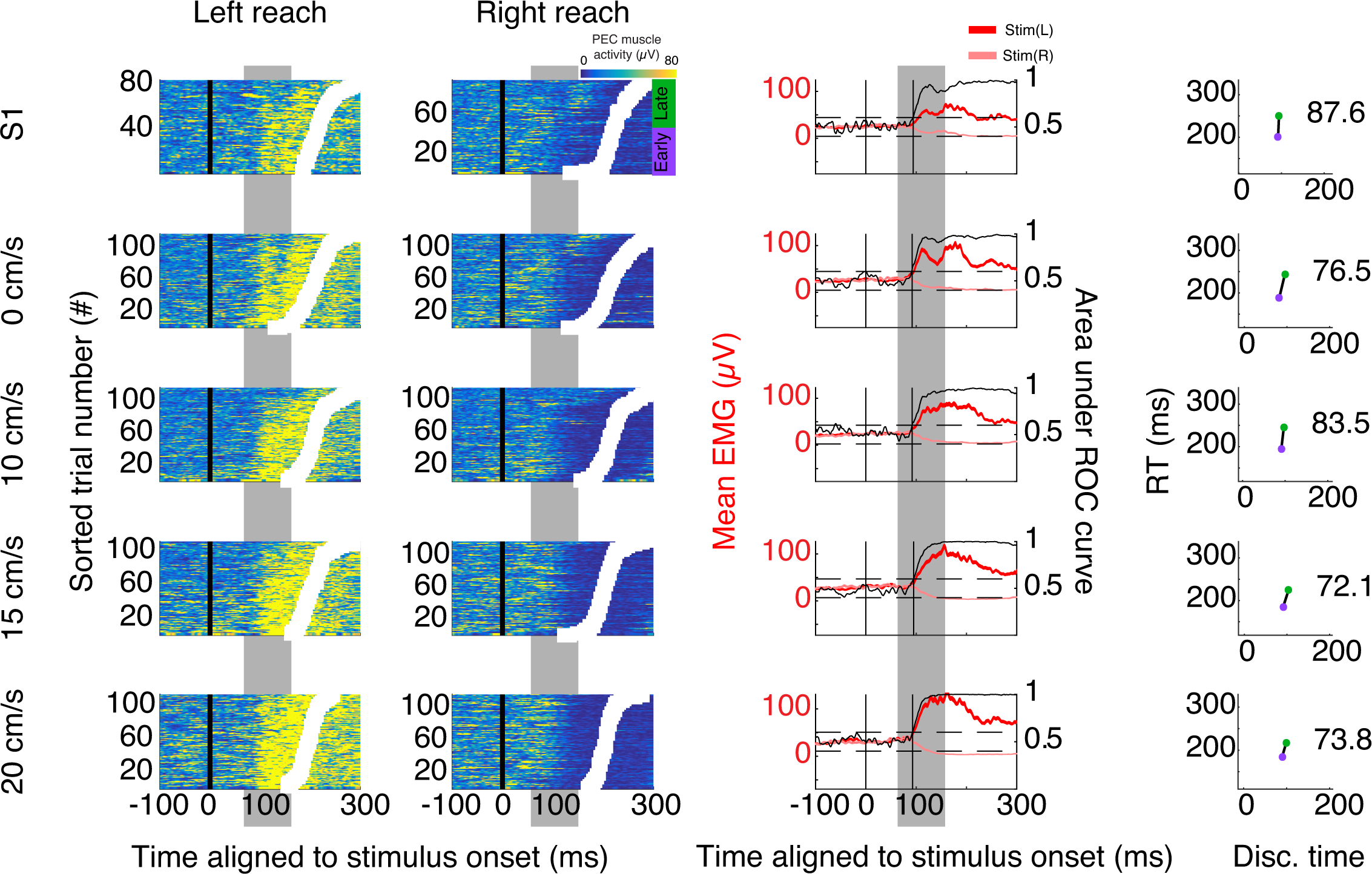
Data from a representative subject in experiment 1, using the same format as in figure 2. From left to right, columns shown trial-by-trial EMG activity for left and right targets sorted by RT, mean +/- SE EMG activity for left and right targets with overlaid time-series ROC, and the RT split analysis for data shown in the corresponding row (the number in each panel is the slope).

In experiment 1, we first quantified SLR prevalence (the proportion of subjects expressing an SLR) towards targets moving at different speeds. Every subject had an SLR in at least one moving target condition (**Figure 3**). We found no difference in SLR prevalence across conditions (10 v. 15 v. 20 cm/s- chi-squared, all p> .05). Recall in experiment 1 that we also presented a static target either in the emerging target paradigm (i.e., the 0 cm/s target, which was intermixed with other moving targets), or in a static task (S1). We found that all 16 subjects (100%) exhibited an SLR to a 0cm/s target, but only 9 of these subjects (∼56%) exhibited an SLR in the standard, static task (S1). This difference in prevalence was statistically significant (chi-squared test: p= 0.00275, χ2= 8.96, df=1), indicating that an emerging target paradigm promotes the expression of an SLR regardless of whether the target is moving or not.

We next examined the latency and magnitude of the SLR. Intriguingly, and reaffirming the impression from the individual data in **Figure** 4, we found no differences in SLR latency across any of the conditions (**Figure 5a;** one-way ANOVA, F(4,53) = .58, *p =* .68). Thus, regardless of target speed, we found that the latency of the SLR remained ∼90 ms. In contrast, we found that target speed impacted SLR magnitude (**Figure 5b)**. Moving from left to right in **Figure 5b**, we found that SLR magnitude increased for faster moving targets (one-way ANOVA across all conditions, F(4, 53)= 15.15, *p=* 2.53e-8; all emerging targets evoked larger magnitude SLRs than static targets; all p<.0005). Indeed, SLRs evoked by the fastest moving target in the emerging target paradigm were approximately twice as large as SLRs evoked by the target in the static paradigm (S1 versus 20 cm/s; unpaired t-test; t= -7.78, p= 3.61e-7, df= 18). Even when removing the influence of both static targets from our analysis (s1 and 0cm/s), we found that faster moving targets in the emerging target paradigm promoted larger magnitude SLRs (one-way ANOVA (10, 15 and 20 cm/s speeds), F(2, 32)= 4.59, *p=* .0183).

**Figure 5.**
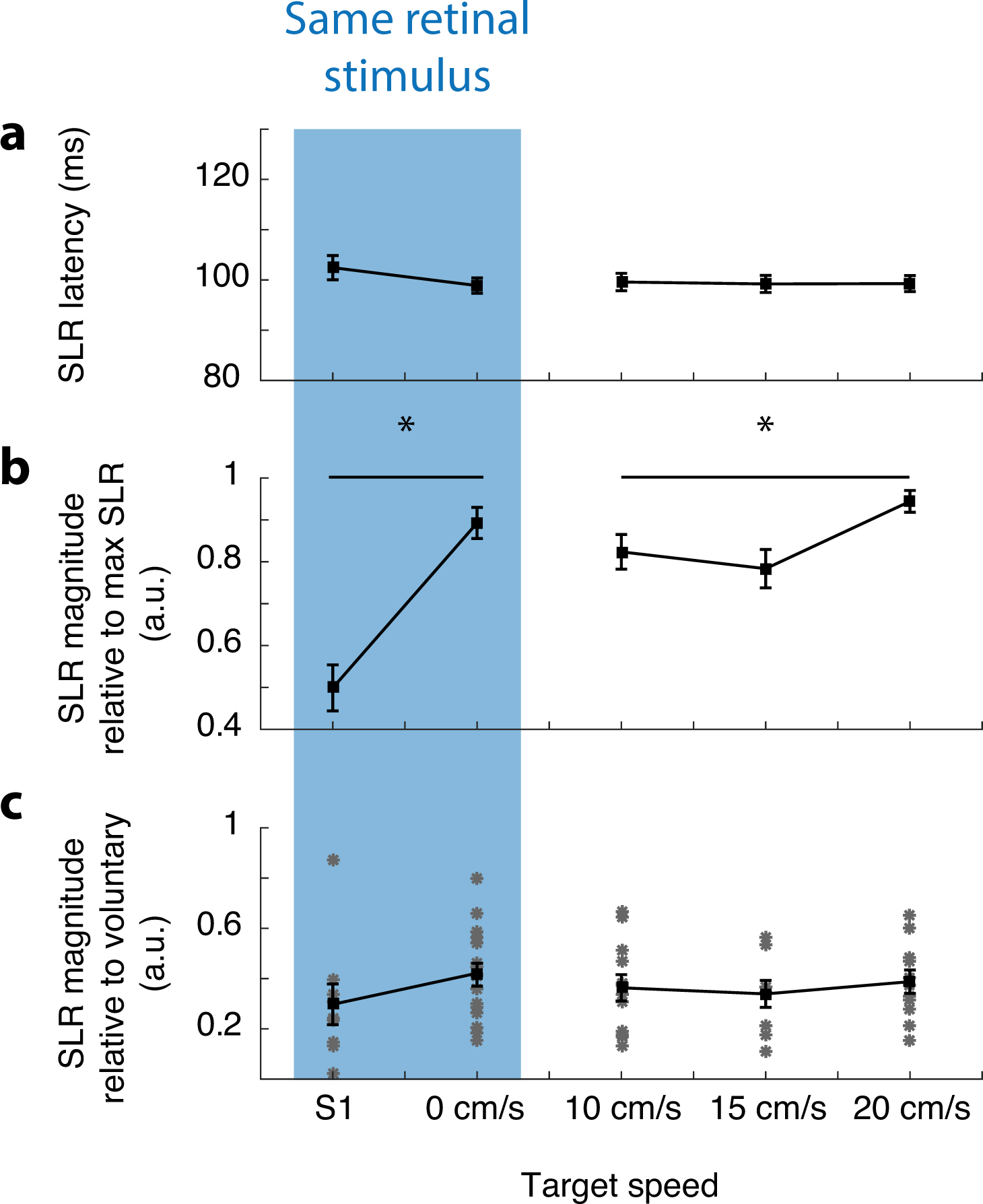
Summary of results from experiment 1, showing (a) SLR latency, (b) SLR magnitude normalized within each subject to the largest SLR across all conditions, and (c) SLR magnitude normalized to the maximum of movement-aligned EMG activity. In c), each asterisk represents data from an individual subject. S1 indicates the results from the static task, which presents the same retinal stimulus to the subject as the 0 cm/s target in the emerging target paradigm. Error bars show SE. * indicates significance (see results text for more details).

In **Figure 5b** (blue rectangle), we noticed differences between the SLR magnitude in the static paradigm (S1) versus that driven by a non-moving (0 cm/s) stimulus in the emerging target paradigm. In these cases, the stimulus presented to the visual system is the same, however SLRs in the emerging target paradigm were almost twice as large (unpaired t-test; t= -6.1, p= 3.02e-6, df= 23). Recall that the SLR prevalence was also higher to the 0 cm/s target in the emerging target paradigm. Thus, depending on the context in which stimuli are presented, they may evoke different SLRs.

Finally, we were interested in how SLR magnitude in the emerging target task related to the maximal movement aligned activity. Previous reports using static tasks observed relatively modest SLR magnitudes which were ∼10-20% as large as muscle activity just prior to movement onset (Pruszynski et al., 2010; Gu et al., 2016; Kozak et al., 2019). As shown in **Figure 5c**, SLRs could be ∼50% (range: 10-90%) as large as the peak of such movement-aligned activity. This is a notable increase in magnitude compared to previous reports. When measured this way, we did not observe any tendency for this scaled measure of SLR magnitude to vary across condition (one-way ANOVA, F(4,53)=.62, *p* = .65).

Overall, targets presented in the emerging target paradigm readily evoked SLRs, regardless of whether they were moving or not. SLRs evoked at similar latencies, however faster moving targets evoked larger magnitude SLRs that occasionally approached levels of movement-aligned activity. We also found that presentation of static targets in the emerging target versus static paradigm evoked larger and more prevalent SLRs, attesting to the influence of task context in favoring SLR expression.

### Experiment 2: High-contrast, fast moving targets evoke shorter latency, larger magnitude SLRs

The most intriguing finding in experiment 1 was that all targets evoked similar latency SLRs. Conversely, all previous reports manipulating visual features found that larger SLR magnitudes were associated with shorter latencies (Wood et al., 2015; Kozak et al., 2019). Lower contrast targets evoke longer latency SLRs (Wood et al., 2015). Thus, to further investigate the relationship between SLR latency and magnitude, in experiment 2 we examined the SLR in response to high and low contrast targets moving at a ‘fast’ (20 cm/s) or ‘slow’ (10 cm/s) speed.

Data from a representative subject in experiment 2 is shown in **Figure 6** (same format as **Figure 4**). This figure illustrates EMG data from both static and emerging target paradigms (compare first two rows to final four rows). In this subject, SLRs in the emerging target paradigm evolved at shorter latencies and attained larger magnitudes in response to high contrast targets (e.g., compare the timing and intensity of EMG activity in the grey box in color plots, or the timing and peak of mean EMG activity in the SLR interval in the 3^rd^ column). As expected, in this subject high contrast targets appear to evoke shorter latency and larger magnitude SLRs than low contrast moving targets. We therefore quantified differences in SLR prevalence, latency and magnitude across the population.

**Figure 6.**
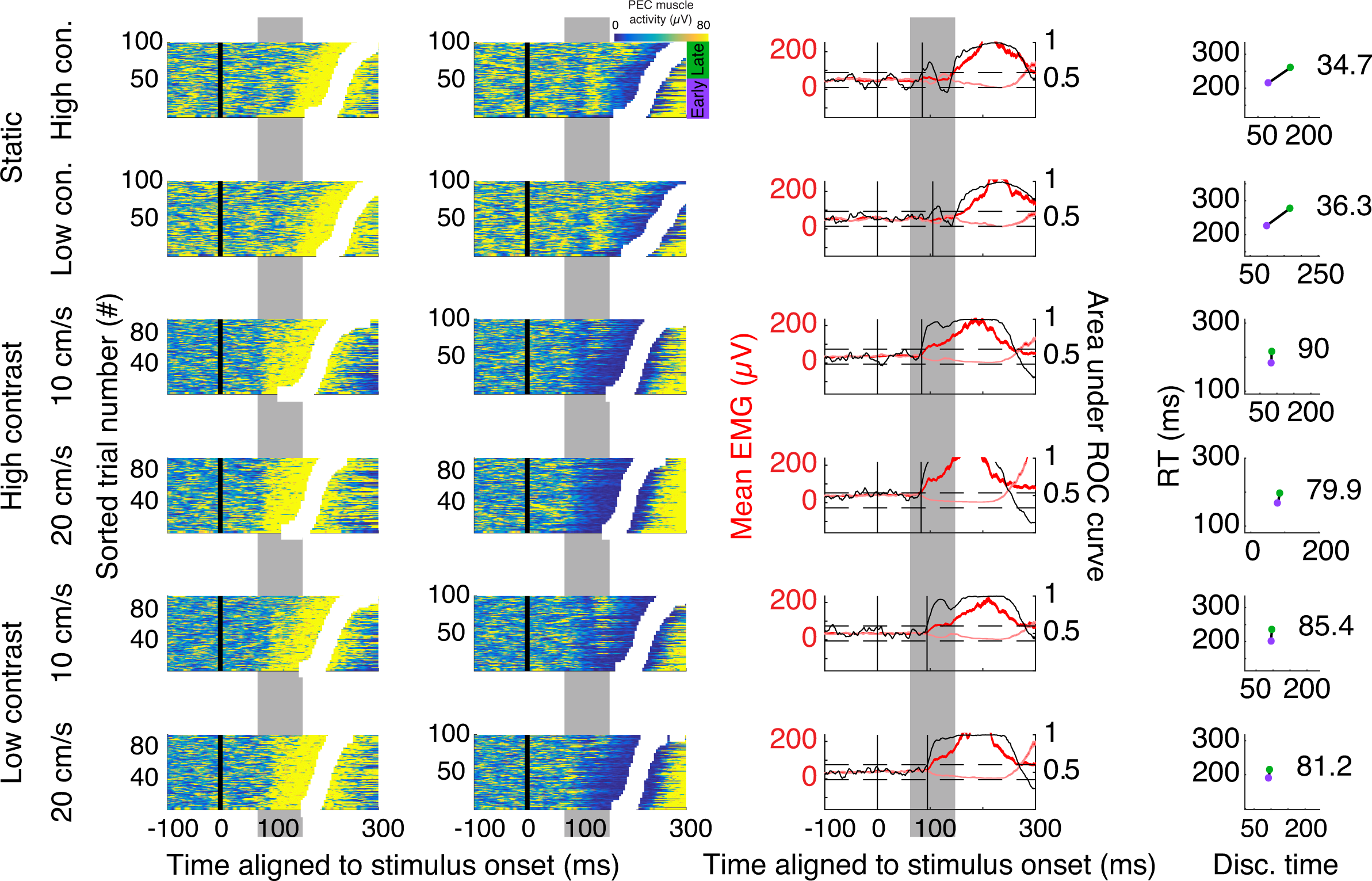
Data from representative subject in experiment 2,using the same format as figure 4.

As in experiment 1, all subjects exhibited an SLR in at least 1 condition (indicated by a diamond above the dotted line in **Figure 3**). In the emerging target paradigm, we found that high contrast targets were more likely to evoke SLRs than low contrast targets (chi-squared test: p= 0.00075, χ2= 11.361, df=1). In fact, low contrast moving targets were no better at evoking SLRs than static targets (slow + fast low contrast moving targets versus high + low contrast static targets: chi-squared test: p>.05, χ2= .056, df=1). Consistent with results from experiment 1, target speed did not impact SLR prevalence (chi-squared, all p>.5).

We next examined SLR latency, and found that target contrast, but not target speed, impacted SLR latency (**Figure 7a**; 2 x 2 ANOVA, *F*(1,47) = 72.85, *p* = 4.1e-11; note that there is no difference within green or blue data points). High contrast targets evoked shorter latency SLRs than low contrast targets (**Figure 7a**; *p* = 2.07e-8, mean within-subject difference ∼14 ms). These results confirm our findings from experiment 1, indicating that target speed does not impact latency.

**Figure 7.**
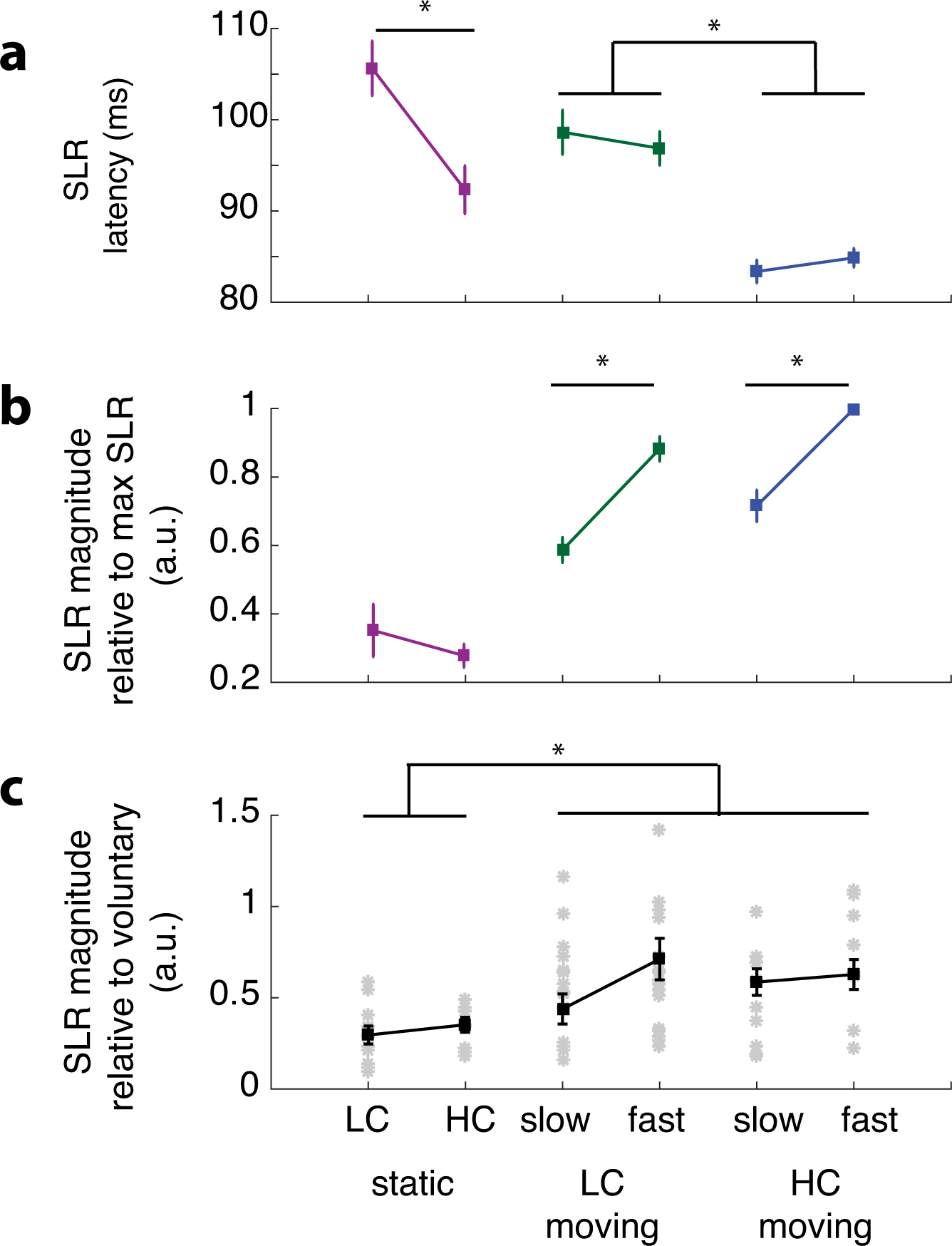
Summary of results from experiment 2 (same format as Fig. 5), with SLR latency (a), SLR magnitude normalized to the within-subject largest SLR (b), and SLR magnitude normalized relative to maximum of movement-aligned activity (c). All values presented as mean +/- standard error. * indicates significance. HC = high contrast. LC = low contrast.

In contrast to experiment 1, we noticed that SLRs evoked in the static task had longer latencies than those in the emerging target paradigm (**Figure 7a;** unpaired t-test; *t*(18) = -3.27, *p* = .004; compare high and low contrast purple data with respective green and blue points). This differed from what we saw in experiment 1, where SLR latencies did not change. We suspect that these differences may relate to the predictability of the time of target appearance (Contemori et al., 2021). Recall that the time of target appearance was fully predictable in experiment 1, but randomized in experiment 2 in order to match a previous study (Wood et al., 2015). These results indicate that emerging targets evoke shorter latency SLRs compared to static targets, if the timing of target appearance is unpredictable.

We next examined the influence of target contrast and speed on SLR magnitude (**Figure 7b**). Here, we found that both target contrast and speed impacted SLR magnitude (2 x 2 ANOVA, speed: F(1,47)=67.8 p = 1.1e-10; contrast: F(1,47)=11.7 p = .001). When matched for speed, higher contrast targets evoked larger magnitude SLRs (**Figure 7b**, all blue points lower than all green points; unpaired t-test; t= 2.7, p= .01, df= 49). We also found that faster moving targets evoked larger magnitude SLRs (unpaired t-test; t= -7.8, p= 4.1e-10, df= 49). This supports the results from experiment 1, where faster moving targets evoked larger magnitude SLRs. We did not observe interaction effects between target speed and contrast (2 x 2 ANOVA, p=.87). We also did not observe any differences in magnitude for static targets.

Next, we compared SLR magnitude to movement-aligned activity (**Figure 7c)** and as seen in experiment 1, observed large variability (**Figure 5c**). However, SLR magnitudes tended to be larger than in experiment 1, reaching ∼75% of movement-aligned activity (∼20% higher than experiment 1; compare **Figures 5c** versus **7c**). In a few cases, SLRs exceeded movement-aligned activity, in the most extreme case approaching 150%. An example of this is shown in **Figure 8**, plotting EMG relative to either stimulus onset (illustrating the SLR; **Figure 8a**) or movement onset (illustrating reach-aligned activity; **Figure 8b**). This figure indicates that recruitment during the SLR interval can exceed movement-aligned activity. **Figure 8a** also provides an example of an SLR that is crisp and distinct from subsequent recruitment, in contrast to the SLRs shown in **Figures 4** and **6**, which blend more smoothly into the ensuing phase of reach-related activity.

**Figure 8.**
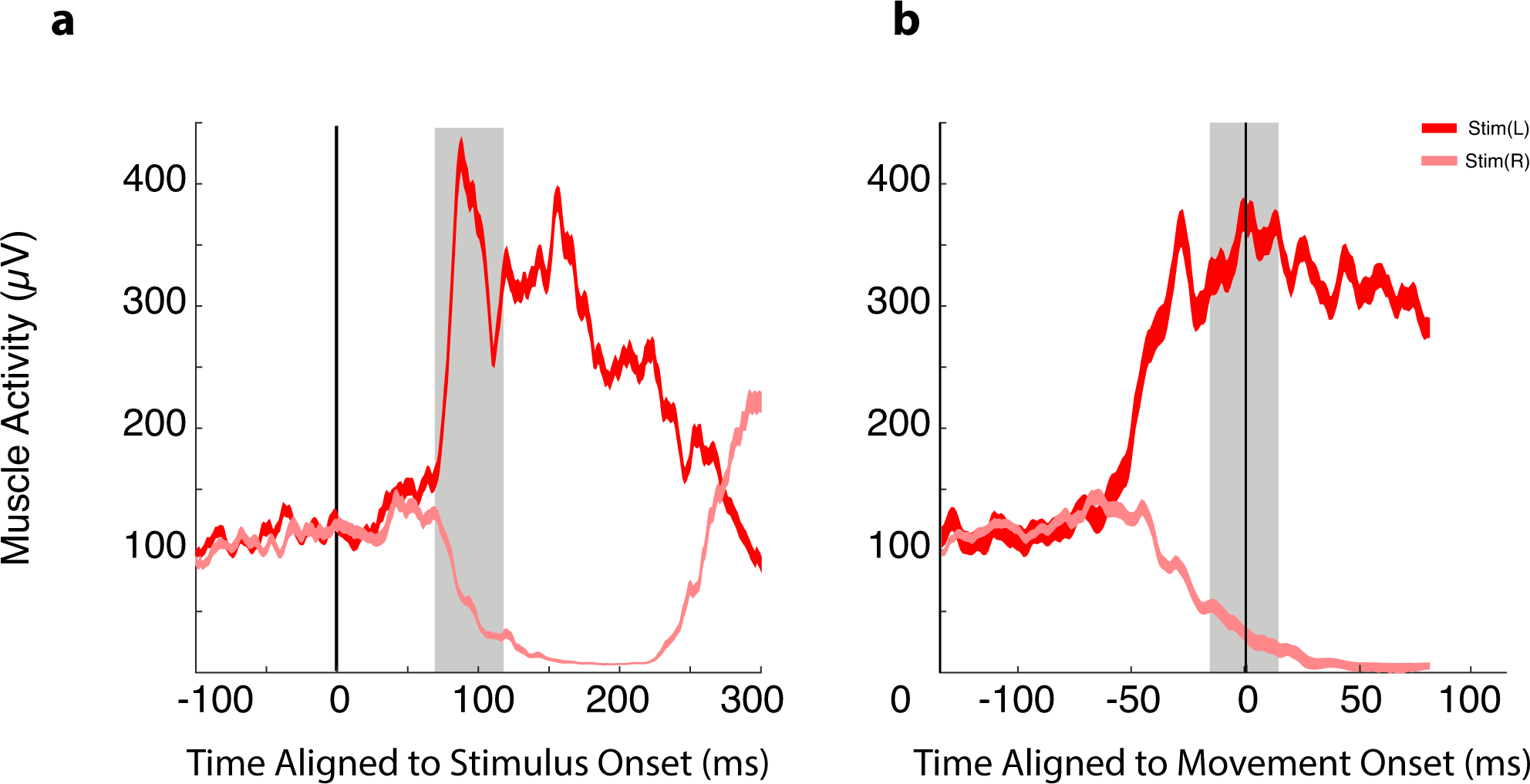
Example subject whose SLR magnitude exceeds the level of EMG activity aligned to target onset. Data, shown as mean +/- SE, are aligned to stimulus onset in a), or movement onset in b). SLR magnitude is measured as area between left and right reaches for the 30 ms interval after the SLR latency (grey box, running from 73-103 ms relative to stimulus onset). For movement-aligned activity, magnitude is measured as area 15 ms before and 15 ms after RT onset (grey box in b).

To summarize, the results from experiment 2 confirmed and extended those from experiment 1, in showing that SLR latency was not influenced by target speed, regardless of target contrast. High contrast, fast moving targets improve SLR expression compared to static targets. Furthermore, we report that SLR magnitude may rival, and even exceed activity attained just before movement onset.

### The SLR and its relationship to reaching behaviour

We further explored the relationship between SLR latency, magnitude, and the ensuing RT. In both experiments, SLR latency and magnitude were negatively correlated with each other (**Figure 9a** exp. 1; R=-.48, p=.00012; **Figure 9d** exp. 2; R=-.41, p=.00038), indicating that larger magnitude SLRs are associated with shorter latency SLRs. Recall that there were no significant latency differences for any target conditions found in experiment 1. This negative correlation indicates that short latency SLRs, even with small variability and a narrow range of latencies (all SLR latencies between ∼90-110 ms **Figure 9a**), still tend to be associated with larger magnitude SLRs.

**Figure 9.**
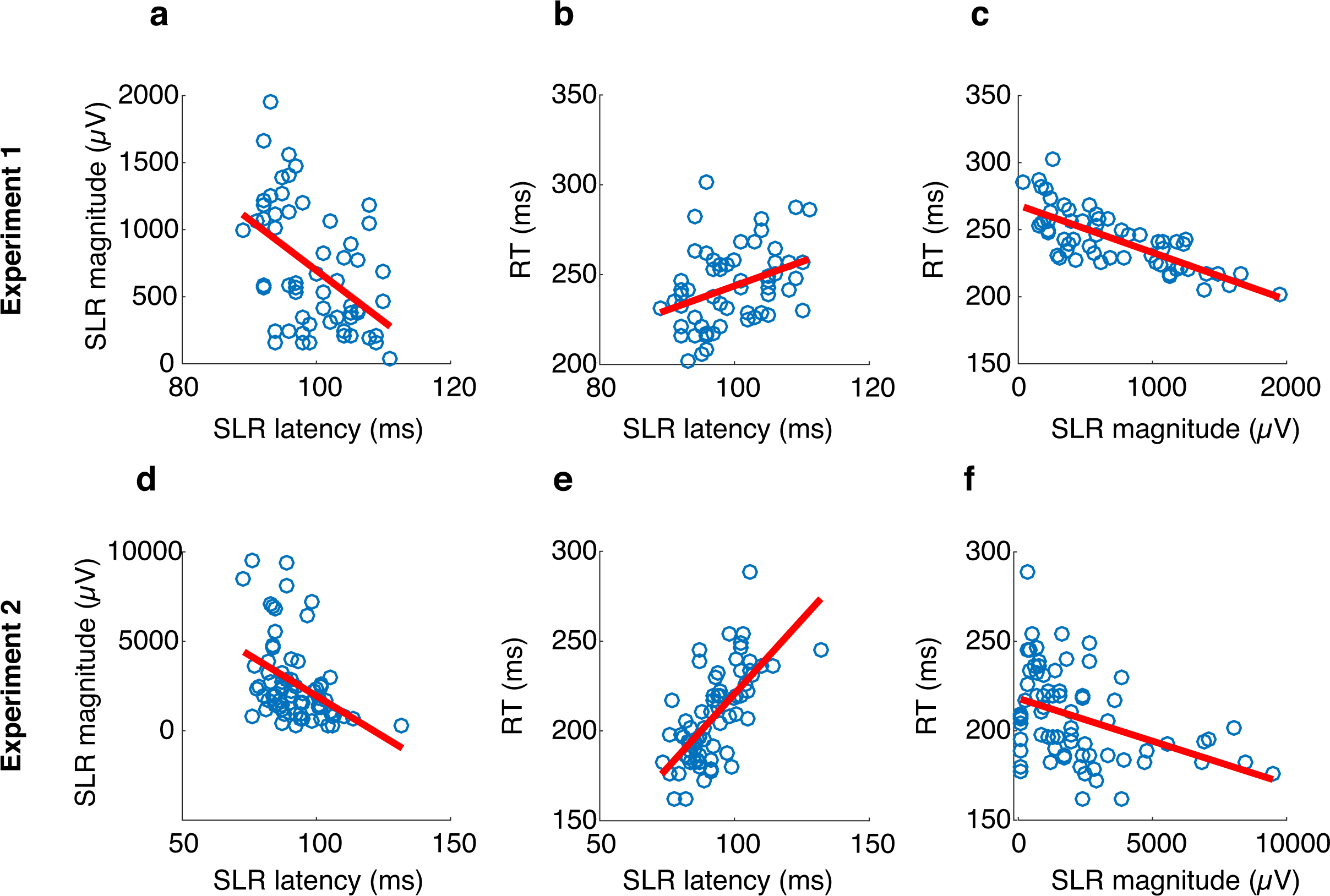
Correlations for SLR latency, magnitude and associated reaction time in experiment 1 (top row) or 2 (bottom row). a, d) SLR latency and magnitude correlations. b, d) SLR latency and RT correlations. c, f) SLR magnitude and RT correlations. All correlations shown are significant. Each blue circle represents data from an SLR+ subject in a unique experimental condition. Solid red lines indicate first order fit of the data.

When examining how the SLR relates to RT, we found a positive correlation between latency and RT in both experiments (experiment 1- **Figure 9b**; R=.36, p=.0047; experiment 2- **Figure 9e**; R=.67, p=8.4e-11); shorter latency SLRs were correlated with more rapid RTs. Conversely, there was a strong negative correlation between SLR magnitude and RT for both experiments (experiment 1- **Figure 9c**; R=-.75, p=8.69e-12; experiment 2- **Figure 9f**; R=-.40, p=.0005), indicating that large magnitude SLRs were associated with more rapid RTs. Thus, robust SLRs are correlated with expedited RTs.

Across both experiments, we found that faster moving (experiment 1) or fast moving high contrast targets (experiment 2) were associated with more rapid RTs (see **Figure 10**). In experiment 1, target speed influenced RTs (one-way ANOVA, F(4, 79)= 2.71, *p=* .036), with RTs decreasing by ∼20 ms (static versus fastest; 250ms+/-25 versus 231ms+/- 16; paired t-test; t= 3.25, p=.005, df= 15). This trend may also be seen in **Figure 4**, which demonstrates a leftward shift of RTs when moving from upper to lower color plots. In experiment 2, both target speed (two-by-two ANOVA, F(1, 107)= 17.33, *p=* 6.5e-5) and contrast (two-by-two ANOVA, F(2, 107)= 103.07, *p=* 3.25e-25) influenced RTs. RTs decreased by ∼62 ms for high contrast targets (static versus fastest; 243ms+/-26 versus 181ms+/- 9; paired t-test; t= 10.2, p=5.7e-9, df= 17) or ∼57 ms for low contrast targets (static versus fastest; 253ms+/-26 versus 196ms+/- 11; paired t-test; t= 9.7, p=2.2e-8, df= 17). Importantly, changes in RTs (**Figure 10**) resembled changes in the latency and magnitude of the SLR (e.g., compare Figure 10a to Figure 5, and Figure 10b to Figure 7). These results indicate a strong relationship between robust SLRs and rapid RTs.

**Figure 10.**
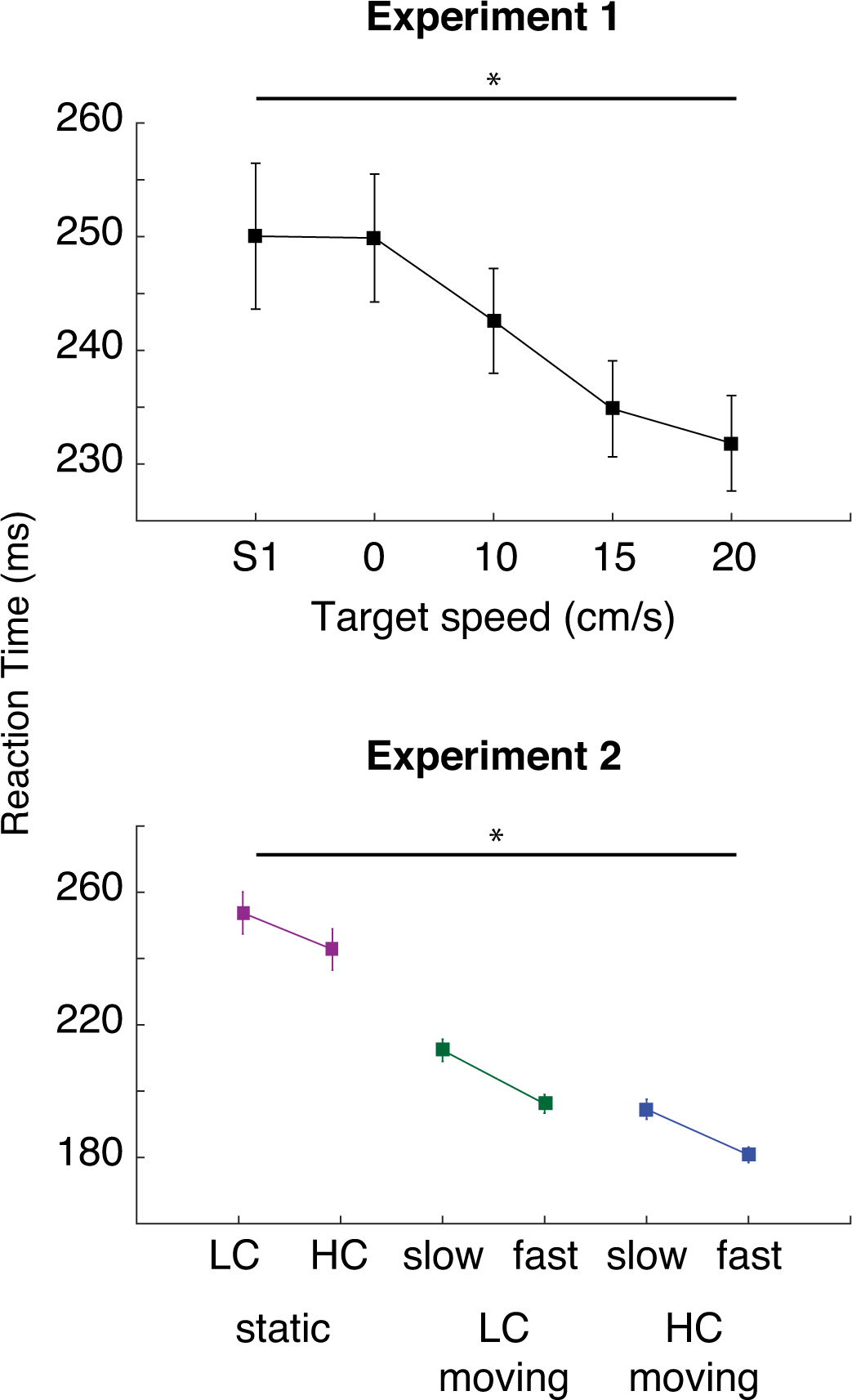
Summary of RTs in experiment 1 (a) and 2 (b), following the formats in figures 5 and 7.

When further examining **Figure 10**, we noticed RT differences between experiments 1 and 2 (∼50 ms), despite both using ‘high contrast’ fast moving targets. These same differences were present in the SLR (**Figure 5a** and **Figure 7a**; unpaired t-test; t= 7.99, p= 2.35e-8, df= 25; ∼15 ms). Recall that experiment 1 and 2 were performed in different apparatuses, with different monitors. We suspect he ‘high contrast’ target in experiment 1 is similar to the low contrast target in experiment 2 (see discussion). In support of this, we found no difference between latencies for low contrast targets in experiment 2 and latencies in experiment 1 (unpaired t-test; t= 1.2, p= .22, df= 25).

Recall in experiment 1 that we presented two different static conditions, one in the standard static task (S1), and one in the context of an emerging target paradigm (0 cm/s) which produced larger and more prevalent SLRs. When examining RTs between these conditions, we observed similar RTs in both tasks (250 +/- 22 ms versus 250 +/- 25 ms; n.s.). Thus, despite differences in SLRs, we observed no differences in RTs. Given the correlation of SLRs to rapid RTs (see above), this result indicates that factors which impact the SLR do not always impact RT in the same way.

To summarize, robust SLRs are correlated with rapid RTs. We found that RTs toward high contrast, fast moving targets decreased by as much as 62 ms. However, we observed conditions where equivalent SLRs preceded different RTs, as well as conditions where different SLRs preceded equivalent RTs. Thus, although robust SLRs promote shorter-latency RTs, RTs are also influenced by periods of muscle recruitment after the SLR.

## Discussion

The brain has a remarkable capacity to rapidly transform vision into action, allowing us to initiate reaches towards a target within fractions of a second. Previous research has shown that some visual stimuli, such those composed of lower spatial frequencies or higher contrast, elicit earlier reaching responses than others (Veerman et al., 2008; Wood et al., 2015; Kozak et al., 2019). Here, by using a new emerging target paradigm (Kozak et al., 2020), we show that high-contrast, moving stimuli are particularly capable of eliciting fast visuomotor responses such as SLRs and expedited reach reaction times. Further, since subjects performed both the emerging target paradigm and a visually-guided reach paradigm used previously, we can better understand the sensory and cognitive factors promoting fast visuomotor responses. Finally, by studying a variety of stimuli in the emerging target paradigm, we can gain further insights into the relationship between the earliest phase of stimulus-related EMG recruitment, subsequent phases of EMG recruitment leading up to movement onset, and the RT of the movement itself. Together, our results contribute to a growing understanding of the interplay between the cortical and subcortical substrates mediating visually-guided reaches in conditions requiring expedited responses.

### Comparison to past studies

Previous research on rapid visually-guided reaches have typically used either target-jump paradigm where a visual target is displaced during an arm movement (Soechting and Lacquaniti, 1983; Day and Lyon, 2000), or a static target reaching paradigm where a suddenly appearing target evokes reaches from a stationary arm position (Pruszynski et al., 2010). While useful, such approaches have limitations: the response of interest evolves during an on-going movement in target-jump experiments, or not all subjects exhibit SLRs in a typical static target paradigm even with intramuscular electrodes (Pruszynski et al., 2010; Gu et al., 2016). In contrast, in the emerging target paradigm, the arm is stationary at the time of target emergence and prevalent fast visuomotor responses can be recorded even with surface electrodes (Contemori et al., 2021; Kozak et al., 2020).

The results in experiment 1 show that larger SLRs are provoked by the appearance of a stationary target in the emerging target paradigm rather than the static task (blue rectangle, **Figure 5**, ∼2 times larger). Larger magnitude SLRs likely relate to the increased proportion of detectable SLRs in the emerging target paradigm rather than static task (100% vs. 56%). We speculate that the increase in SLR magnitude and prevalence in the emerging target paradigm is due to a combination of a signal of implied motion of the target behind the barrier (Kozak et al., 2020) and certainty of target appearance (Contemori et al. 2021). Implied motion is critical, as a recent study presenting only moving targets did not report SLRs (Lara et al., 2018). Further, temporally predictable targets still have to be presented within the emerging target paradigm, as smaller magnitude and less prevalent SLRs are elicited in the static tasks used here despite a predictable gap between the offset of a central hold stimulus and target presentation (Pruszynski et al., 2010; Wood et al., 2015).

Past work has shown that stimuli composed of progressively higher contrast (Wood et al., 2015) or lower spatial frequencies (Kozak et al., 2019) elicit progressively larger magnitude and lower latency SLRs. Somewhat surprisingly, manipulating stimulus velocity did not elicit this inverse relationship between SLR latency and magnitude. Instead, SLR latency remained invariant while SLR magnitude was greatest for the fastest moving stimulus; this was observed both in experiment 1 (**Figure 5**) and in experiment 2 for stimuli of similar contrast (**Figure 7**). This finding demonstrates that the latency and magnitude of the SLR are dissociable. We speculate that this dissociation relates to the displacement of the stimulus across sequential display frames, with larger displacements increasing the vigor of sensory transients that remain timed to the onset of the first frame. Future studies should examine the impact of subsequent display frames on the SLR, given that Contemori and colleagues (2021) found that a stimulus flashed for a single frame (∼16 ms) provokes earlier and larger SLRs than moving targets; it is possible that flashed stimuli may provoke particularly vigorous sensory transients from the retina.

Our observations of scaled EMG responses for faster moving targets resembles the phenomenon of speed coupling in the interception timing literature, where participants adjust hand velocity to match target velocity (Savelsbergh et al., 1992; van Donkelaar et al., 1992; Carnahan and McFadyen, 1996; Brouwer et al., 2000, 2002, 2003). Speed coupling may reflect the temporal accuracy required to intercept targets moving at different speeds, rather than direct coupling to the speed of the target (Brouwer et al., 2000, 2002; Tresilian and Lonergan, 2002; Caljouw et al., 2004). Aspects of our results may be consistent with speed coupling, particularly since target motion before disappearance cues the subject about upcoming target velocity. On the other hand, the extremely short latency of the SLR, regardless of target speed, is consistent with it arising from a feed-forward response. Future work should examine SLR magnitude across target velocity in the absence of any predictability, and compare such SLRs to subsequent recruitment preceding the RT.

### Relating the SLR to subsequent RTs

SLRs represent the first change in muscle recruitment following visual stimulus presentation. As such, they demarcate the completion of the most rapid sensory-to-motor transformation for visually-guided reaching. Past reports show that larger magnitude SLRs precede shorter latency movements on a trial-by-trial basis (Pruszynski et al., 2010; Gu et al., 2016; Contemori et al., 2021), and here we show, when pooling across all conditions in which subjects exhibited an SLR, that larger magnitude and earlier SLRs precede shorter reach RTs (**Figure 9;** see also Kozak et al., 2019). These correlative relationships, are consistent with a functional contribution of the SLR to the forces necessary to overcome the limb’s inertia. However, the SLR is often followed by sustained levels of muscle recruitment preceding the RT. Consequently, muscle recruitment during the SLR interval is not fully predictive of the RT. Nearly identical SLRs can precede different RTs (e.g., in Experiment 1, different RTs toward 0 deg/s and 20 deg/s stimuli despite similar SLRs), or different SLRs can precede the same RTs (e.g., in Experiment 2, similar RTs toward fast moving low-contrast or slow moving high-contrast stimuli despite different SLRs). These distinctions between SLRs and RTs attest to continued recruitment after the SLR, and the contribution of such recruitment to overcoming the limb’s inertia. Judicious selection of stimulus parameters, e.g., to elicit robust SLRs but longer RTs, may aid the study of the fast visuomotor responses by reducing the overlap between the SLR and activity aligned to movement onset.

Previous reports of SLRs emphasized their relatively small magnitude, reaching ∼10-20% of the volitional recruitment achieved before movement onset. Such modest levels of recruitment during the SLR interval can induce subtle motion toward the target, even if a subsequent movement is withheld or proceeds in a different direction (Gu et al., 2016; Atsma et al., 2018). A remarkable aspect of the current results is that we occasionally observed SLR recruitment that equaled if not exceeded volitional, movement-related activity (**Figure 8**). As mentioned above, increased SLRs magnitude likely relates to the increased propensity of detecting an SLR. Further, for many subjects, the shortest RTs occurred essentially right after the SLR interval (e.g., see shortest RT trials for the 20 cm/s targets in **Figures 4** and **6;** trials in lowest rows). In such trials, EMG recruitment during the SLR interval is clearly sufficient to overcome the limb’s inertia to produce a detectable RT. On such trials, any distinction between the SLR and subsequent volitional recruitment is blurred; phases of recruitment that are separated for longer- RT movements are essentially indistinguishable for shorter-RT movements. We are struck by the resemblance of the single-trial EMG data shown in **Figures 2, 4, and 6** to raster plots of activity of the superior colliculus during express and non-express saccades (Edelman and Keller, 1996; Dorris et al., 1997; Sparks et al., 2000). For shorter latency movements, be they express saccades or short-RT movements in the emerging target paradigm, phases of recruitment tied to stimulus onset merge into those phases of recruitment loosely termed ‘volitional’ for longer-latency movements.

### Plausible neural substrates

The superior colliculus (SC) has been implicated as a potential candidate mediating the SLR (Corneil et al., 2004; Pruszynski et al., 2010). The SC is a midbrain structure whose intermediate and deep layers (SCi) interface with premotor structures for eye, head, and upper limb movements within the reticular formation (Corneil and Munoz, 2014), whereas superficial layers respond to visual input. Our results add further circumstantial evidence for a role for the SC in the SLR. Previous work established that both the SLR and visual responses in the SC preferentially respond to low spatial frequency (Chen et al., 2018; Kozak et al., 2019) and high contrast targets (Li and Basso, 2008; Marino et al., 2015; Wood et al., 2015). Moving targets also evoke robust SC responses (Marrocco and Li, 1977), and we have shown here that moving targets also evoke robust SLRs. Importantly, the target speeds used in the current study resemble those used in the Marrocco and Li (1977) study, which evoked robust responses in the intermediate and deep layers of the SC where visuomotor and motor neurons activated before arm movements reside (Werner et al., 1997; Stuphorn et al., 1999; Philipp and Hoffmann, 2014). The hypothesis that the SC mediates the SLR requires further testing with casual manipulation in tasks that promote SLR expression. Work reporting that SCi inactivation impacted the selection of reach targets but not reach kinematics (Song et al., 2011) employed paradigms that would not promote SLR expression.

An open question is how visual input from target motion accesses the SC, and how features of the emerging target task potentiate SLRs. Visual information may access the superficial SC directly via a retinotectal route (summarized in: (May, 2006)), or indirectly via a retino-geniculo-striate loop through cortex, and then on to the SC (Schiller et al., 1979). The SC also receives projections from higher-order cortical centers such as MT ((Fries, 1984; Nhan and Callaway, 2012); an area which responds to target motion). As MT response latencies to visual stimuli are ∼60-70 ms in NHPs (Schmolesky et al., 1998; Smith et al., 2005), which are concurrent with SLR latencies on NHP neck muscles (Corneil et al., 2004), motion processing in MT may not be directly involved in relaying visual transients that evoke the SLR. Instead, as implied motion strongly activates MT (Krekelberg et al., 2005), top-down signals from MT may strongly activate pre-target activity in the SC, increasing the vigor of the SC’s response when the target appears beneath the occluder, and subsequently increasing the magnitude of the SLR.

### Methodological considerations

All of our subjects exhibited an SLR in at least one stimulus condition. This replicates the high prevalence of SLR detection in the emerging target paradigm (Kozak et al., 2020; Contemori et al., 2021), and affirms the contention that most if not all subjects are capable of exhibiting SLRs in an appropriate paradigm. However, subjects did not necessarily exhibit an SLR to every stimulus in the emerging target paradigm, even for high-contrast, fast moving stimuli (**Figure 2**). We speculate that this result may be due, at least in part, to how we detect the presence of the SLR with the RT split analysis, which allows for a direct comparison of SLR prevalence to those reported elsewhere which also used this method (e.g.,(Kozak et al., 2019, 2020; Contemori et al., 2021)). The RT split analysis is somewhat conservative in that it requires an SLR to both short- and long-latency trials. SLR- observations could therefore include those situations where an SLR precedes the short- but not long-RT trials (see top two rows of **Figure 4**, which appear to exhibit SLRs for the shorter-than-average RTs). The RT split analysis is also more suited to paradigms engendering a larger degree of RT variance. As noted previously (Kozak et al., 2020), one potential complication of the emerging target paradigm is that it may shorten RTs too much, not only reducing RT variance but also leading to a potential overlap of SLR with activity linked to movement onset. As noted above, judicious stimulus selection may help avoid such overlap, but we also expect the continued evolution of SLR detection methods suited to a particular paradigm.

We noted large differences in the latencies of fast visuomotor responses across experiments, with SLRs evolving ∼15 ms earlier and reach RTs being ∼50 ms shorter in experiment 2 compared to experiment 1. The most likely reason for these differences is the contrast of the stimuli, as the VPIXX projector used in experiment 2 has a higher level of contrast (275 cd/m2) than the standard monitor used in experiment 1 (110 cd/m2). Consistent with this, the SLR latencies from the high-contrast target in Experiment 1 (**Figure 5b**) resemble those of the low contrast target in experiment 2 (**Figure 7a**). The large difference in reach RTs across experiments, even to targets associated with similar latency SLRs, may also relate to postural differences in the exoskeleton-based Kinarm robot in experiment 1 versus the endpoint-based Kinarm robot in experiment 2.

## Conclusions

The work presented here demonstrates that all subjects can generate SLRs, but are more likely to do so in an emerging target paradigm. As a consequence, reliable SLRs can be recorded even with surface EMG, and here we use this paradigm to show that targets moving at faster speeds elicit larger magnitude SLRs, whereas higher contrast targets decrease SLR latency. Larger magnitude and shorter-latency SLRs tend to precede expedited RTs. As the SLR is the first wave of muscle recruitment tied to visual target onset, muscle recruitment during this interval may provide a unique window into the contribution of the tecto-reticulospinal signaling that is distinct from subsequent phases of target-related recruitment of corticospinal or cortico-reticulo-spinal circuits. Rather than being directly involved in the SLR, cortical areas such as MT may serve to prime this circuit to produce the first wave of muscle recruitment in situations when time is of the essence.

